# A floor-field model of tau-taxol competition explains tau envelope dynamics on taxol-stabilized microtubules

**DOI:** 10.1101/2024.11.19.624338

**Authors:** Carole Chevalier, Valerie Siahaan, Rahul Grover, Stefan Diez, Ludger Santen

## Abstract

Tau is a microtubule-associated protein (MAP) that is highly expressed in neurons. Recent in-vitro studies have shown that tau molecules can segregate into cohesive ‘envelopes’ on paclitaxel (taxol) stabilized microtubules. These envelopes act as selective barriers gating the motor transport and protecting microtubules against the action of severing enzymes. However, the mechanisms underlying the formation of these envelopes remain unclear. Recent in vitro reconstitution assays revealed that the formation of tau envelopes induces a compaction of the underlying microtubule lattice, whereas taxol binding induces lattice expansion, indicating competitive binding between tau and taxol on microtubules. In this study, we use physical modeling to investigate how this competition between tau and taxol regulates the formation of tau envelopes. Our model includes static and dynamic floor-fields, concepts adapted from pedestrian dynamics. By comparing our simulation results with experimental data, we demonstrate that tau-taxol competition on the microtubule lattice is sufficient to explain the features of the formation and disassembly of tau envelopes. Notably, tau-tau interactions are not necessary to account for the steadiness of these envelopes.

## 1 Introduction

Microtubules serve three essential functions in living cells: they form the structural cytoskeleton network, determining cell shape; they play a crucial role in separating chromosomes during mitosis; and they act as tracks for the intracellular transport of molecules and organelles, which are vital for cell function. Microtubules are dynamic polymers build up of alpha and beta-tubulin that form tubulin dimers. These tubulin dimers assemble head-to-tail into protofilaments that enclose into a hollow tube. Each MT generally consists of 13 protofilaments but typically ranges between 12-15 protofilaments [1, 2]. Microtubules interact with various molecules known as microtubule-associated proteins (MAPs), which play key roles in stabilizing, orienting, and cross-linking microtubules [3, 4]. Investigating how these MAPs influence microtubule functions is a fundamental area of study. Tau, abbreviated from ‘tubulin-associated unit’, was one of the first MAPs to be characterized. It is predominantly found in neuronal axons [5], where it plays a crucial role in maintaining the integrity of the cytoskeleton network [6]. However, tau is also implicated in several pathological situations. Dysfunction of tau proteins is implicated in the development of neurodegenerative conditions commonly referred to as tauopathies, which include Alzheimer’s disease, Pick’s disease and frontotemporal dementia [7, 8]. At the cellular level, studies indicate that elevated levels of tau can alter the distribution of various organelles, suggesting a role for tau in regulating intracellular transport [9, 10].

In this paper, we study a novel aspect of tau’s function: its propensity to assemble into stable clusters termed as ‘tau islands’ or ‘tau envelopes’ along microtubules [11–13]. These tau envelopes act as protective coating, shielding microtubules from severing induced by enzymes such as katanin or spastin and as a selective barrier allowing the processive movement of certain molecular motors while inhibiting others, thus modulating intracellular transport [11, 12]. However, the mechanism underlying tau envelope formation is not known. To understand this mechanism is not only a crucial question in medicine, but also an intriguing topic in fundamental biology and physics.

A first computational study attempted to explain tau envelope formation through tau-tau interactions [14], as suggested in [13]. The result was that several small tau patches were formed but did not assemble into stable envelopes of several micrometers in length. Consequently, a different mechanism must be considered to explain the tau envelope formation as observed in experiments.

In the present study, we consider paclitaxel (taxol), a chemotherapeutic agent commonly used to stabilize microtubules and halt their dynamics during *in vitro* experiments [15–17]. Interestingly, taxol has been shown to interfere with tau binding both *in vivo* [18] and *in vitro* [13]. Specifically, taxol disrupts the formation of tau envelopes, highlighting a competitive interaction between tau and taxol for microtubule binding. Tau envelopes compact the lattice of taxol-stabilized microtubules, in contrast to taxol’s known role in expanding the microtubule lattice [19]. Notably, tau can bind to tubulin dimers already occupied by taxol, effectively displacing taxol from the microtubules [13]. Because taxol binds only transiently to microtubules, it permits lattice compaction upon tau binding.

Consequently, the idea emerge that tau-taxol competition can have an important role in the tau envelope formation and disassembly features observed experimentally. To study this hypothesis, we model the tau-taxol competition using a floor-field approach. The floor-field concept, originally introduced for modeling pedestrian dynamics [20, 21], was inspired by chemotaxis process used by insects and cells for communication [22]. Mammal, insect, or cellular dynamics are often modeled using discrete lattices, where entities move stochastically with a specific stepping rate. These lattices, known as lattice gas or cellular automata, can incorporate a floor-field to introduce variability in stepping rate. This variability can be spatial, where the stepping rate differs across the lattice, a scenario referred to as a *static floor-field*. Alternatively, the variability can be temporal, where the stepping rate changes over time, leading to what is known as a *dynamic floor-field*. In pedestrian kinetics, spatial rate variations can model obstacles like walls, while temporal rate variations model virtual traces of pedestrians which influence the motion of other individuals [23]. Here, we apply a dynamic floor-field to account for the influence of tau and taxol molecules adsorbed on the microtubule, on the dynamics of tau envelope formation. Additionally, we introduce irregularities on the lattice modeled by a static floor-field, to account for the observed halting of envelope spreading in experiments [12]. Moreover, we include a hysteresis loop allowing for the differences in tau densities observed during tau envelope formation and disassembly. By comparing the experimental measurements previously presented in [12] to the model results, we show that tau-taxol competition, with the associated local microtubule lattice compaction-expansion alone can explain the tau envelope formation without the need of additional tau-tau interactions.

## 2 Definition of the floor-field model

### 2.1 Microtubule as 1D lattice

To investigate the effect of tau-taxol competition on tau envelope formation on microtubules, we use a coarse-grained model of adsorption and desorption of tau and taxol on a one dimensional lattice with a floor-field as illustrated in the Fig. 1. In this model, one site corresponds to four tubulin dimers, which is the number of tubulin dimers covered by a single tau molecule. As microtubules generally consist of 13 protofilaments, we allow 13 tau molecules to bind to one site. As one taxol can attach on each tubulin dimer, we allow 4 *×* 13 = 52 taxols on each site. Both tau and taxol have their *intrinsic* adsorption (desorption) rates which are respectively 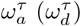 and 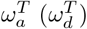. The tau molecules can diffuse along the lattice in both directions with a diffusion rate *s* whereas the taxol molecules do not diffuse. The stationary density of such a model is well known and called the Langmuir density [24]:

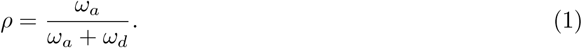

**Figure 1:**
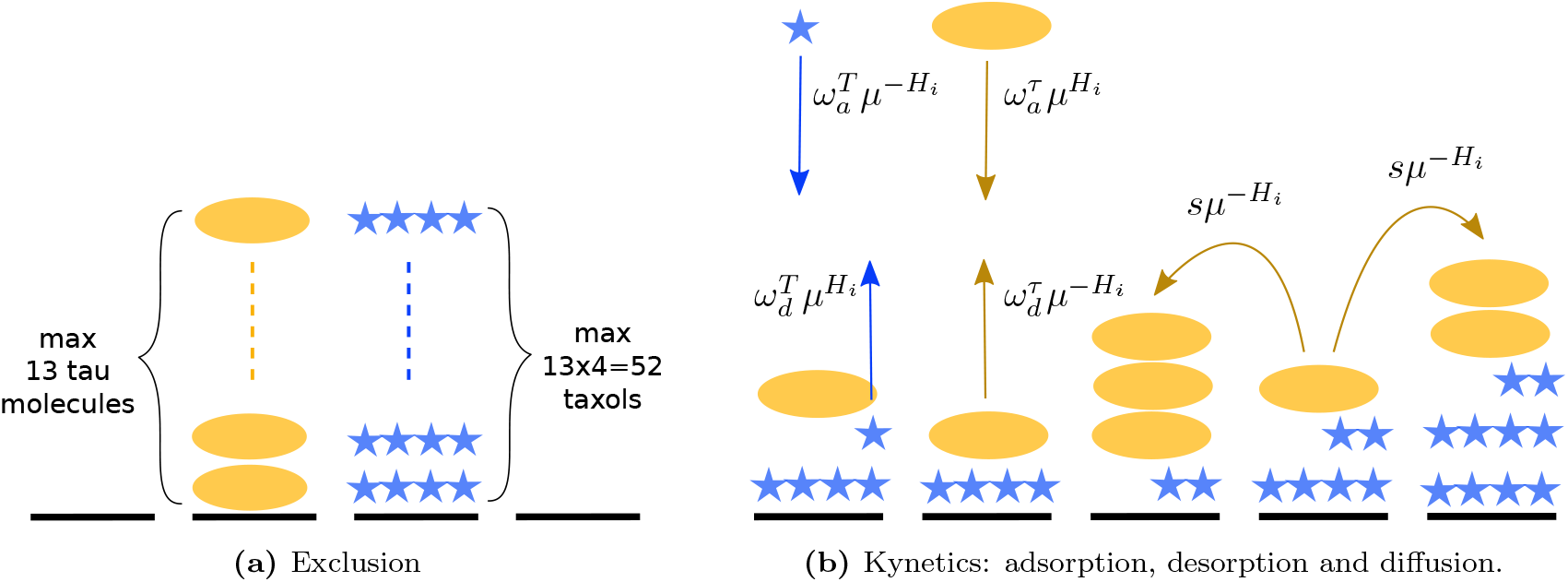
Schematics of the physical model. Microtubule lattice sites are represented by black dashes, with yellow ovals representing taus and blue stars representing taxols. (a) Exclusion is considered such as the maximal number of tau on one site is 13 and the maximal number of taxol allowed on one site is 52. (b) Kinetics of tau (taxol) on the lattice governed by the tau (taxol) adsorption rate 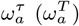, the tau (taxol) desorption rate 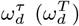, the tau diffusion rate *s*, the affinity modification factor *μ* and the floor-field *H*_*i*_.

Each tau and taxol has its own Langmuir density depending on its respective adsorption and desorption rates. For completeness, we note that the diffusion rate *s* can only influence particle density if it is of the same order as *ω*_*a*_ and *ω*_*d*_, which is not the case for tau molecules on microtubules [12]. The diffusion rate *s* tunes the homogeneity of the tau density along the microtubule.

### 2.2 A dynamic floor-field to model the tau-taxol competition

Using a floor-field, we model the tau-taxol competition and the influence of the microtubule lattice compaction. We do not model the states of the tubulin dimers (compacted or expanded), but only the resulting self-attractive phenomenon induced by tau on microtubules. Specifically, by compacting the microtubule lattice, the tau molecules already adsorbed on the microtubule attract additional tau molecules to adsorb on the microtubule. Since we account for the 13 protofilaments by allowing 13 taus at each site, the increase in floor-field with the number of tau adsorbed on the lattice can be seen as a transversal compaction of the microtubule. This approach effectively represents the tau patches formation as envelopes. Similarly, we model the repulsive effect of taxol on tau adsorption (by expanding the microtubule lattice) and conversely the repulsive effect of tau on taxol adsorption (by microtubule lattice compaction). To allow the spreading of the tau envelopes, we consider that the adsorbed molecules at each site influence the kinetic adsorption/desorption rates on their two nearest neighboring sites.

We introduce a dynamic, local floor-field *H*_*i*_, that varies based on the molecules adsorbed on the microtubule. Specifically, it depends on the local (on a site *i*) numbers of tau and taxol, and on the first neighboring numbers of tau and taxols (on sites *i* − 1 and *i* + 1) to consider an influence of the first neighboring molecules. The term ‘dynamic’ refers to the temporal variation of the floor-field as it changes with the fluctuation in the number of adsorbed molecules on the microtubule. The expression of the floor-field is given by:

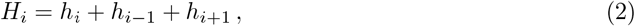

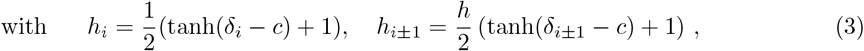

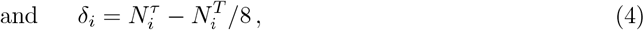

where 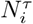 is the number of tau molecules on site *i*, 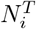 is the number of taxols on site *i* and *h* is called the coupling factor. In order to have envelopes of only tau molecules and not envelopes of taxols, we consider a less important influence of taxol on the floor-field by dividing its number by eight in *δ*_*i*_. Since there are four times as many taxols allowed on a site as taus, the influence of all taxols is half that of taus. Therefore, 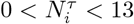 and 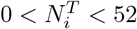 which leads to −6.5 *< δ*_*i*_ *<* 13.

The experimental results on tau envelopes have been described as ‘kinetically distinct phases of tau on microtubules’ [12]. The first phase consists of high-density patches of slow-diffusing tau, the envelopes, while the second phase, the envelope surroundings, is characterized by fast-diffusing tau at low density. Hereafter, we will refer to this second phase simply as ‘the surroundings’.

We choose a hyperbolic tangent shape of the floor-field to account for a sharp transition between the two phases (Fig. 2). A site *i* has a floor-field effect *h*_*i*_ at the site *i, h*_*i*−1_ at its right neighbor site *i* − 1, and *h*_*i*+1_ at its left neighbor site *i* + 1. Each hyperbolic tangent has a value between 0 and 1 by dividing the function by 2 and adding 0.5. Consequently, the site *i* produces a floor-field between 0 and 1, the sites *i* − 1 and *i* + 1 produce a floor-field between 0 and *h*, and the total floor-field *H*_*i*_ has a value between 0 and 2*h* + 1. The coupling constant *h* tunes the strength of the floor-field due to the neighboring sites (*i* − 1 and *i* + 1). A large *h* means a high floor-field at the neighboring sites and therefore a fast envelope growth.

**Figure 2:**
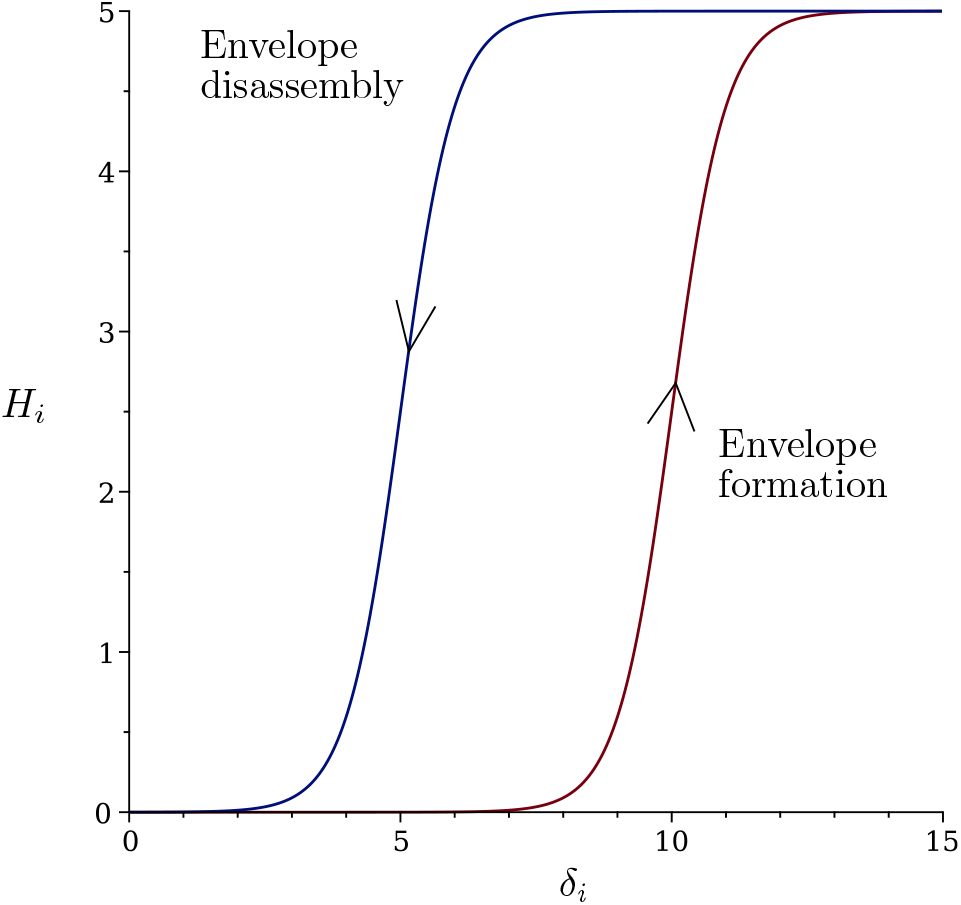
Graph of the floor-field function as define in Eq. (2) for *c*_*f*_ = 10 (red) and *c*_*d*_ = 5 (blue). To give a simple and meaningful illustration we consider *δ*_*i*−1_ = *δ*_*i*_ = *δ*_*i*+1_. The coupling factor is fixed at *h* = 2. Together, the two *H*_*i*_ curves form a hysteresis loop with ∆*c* = 5.

The floor-field represents a local phase transition depending on the difference between the number of tau and taxol molecules adsorbed on the site *i*, denoted as *δ*_*i*_, as well as the corresponding differences at the first neighboring sites *δ*_*i*−1_ and *δ*_*i*+1_. Since −6.5 *< δ*_*i*_ *<* 13 for all *i*, the center of the hyperbolic, denoted as *c*_*f*_, must lie within the specified interval but be sufficiently distant from the boundaries to allow for a phase transition. The parameter *c*_*f*_ determines the specific tau and taxol densities (i.e. *δ*_*i*_, *δ*_*i±*1_), at which the phase transition occurs. Consequently, *c*_*f*_ also tunes the tau and taxol densities (i.e. *δ*_*i*_, *δ*_*i±*1_) allowed in each phase.

Moreover, *c*_*f*_ also tunes the rate of tau envelope nucleation dependent on the stationary densities. When *c*_*f*_ is close to the value of *δ*_*i*_ given by the stationary densities of tau and taxol in the surroundings, the envelopes are more likely to nucleate, as the conditions for *δ*_*i*_ and *δ*_*i±*1_ leading to the envelope formation are frequently met. Conversely, if *c*_*f*_ is far from the value of *δ*_*i*_ given by the stationary densities in surroundings, envelopes nucleation is rare as the *δ*_*i*_ and *δ*_*i±*1_ required for envelope formation are seldom achieved. Notably, the number of adsorbed taxol molecules decreases *δ*_*i*_ and *δ*_*i±*1_, shifting it away from *c*_*f*_, which explains the reduced envelope coverage observed on the microtubules stabilized with taxol at low tau concentrations. The value of *H*_*i*_ influences the adsorption and desorption rates of tau and taxol and the diffusion rate of tau as follows:

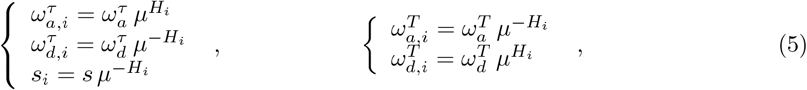

were *μ* ≥ 1 is called affinity modification factor. When *H*_*i*_ = 0, implying no floor-field on site *i*, the rates are not modified since *μ*^0^ = 1. However, when *H*_*i*_ *>* 0, *μ* calibrates the amplitude of the floor-field on the rates. Consequently, *μ* tunes the stationary density in the envelopes, by modifying the effective adsorption and desorption rates, and its homogeneity, by modifying the effective diffusion rate. The dependency of each rate on the floor-field is chosen to reflect specific interactions: tau affinity is increased at higher concentration of tau and lower concentration of taxol on the lattice. Oppositely, taxol affinity is increased at lower concentration of tau and higher concentration of taxol on the lattice. Additionally, tau diffusion increases as tau concentration decreases and as taxol concentration increases. In particular, this means that the affinity of tau to a site *i* is enhanced if there are enough taus comparing to taxols on this site *i* (*δ*_*i*_ *>* 0) or on the left neighbor site (*δ*_*i*−1_ *>* 0), or on the right neighbor site (*δ*_*i*+1_ *>* 0). This models the increase of tau affinity due to the presence of other tau molecules that compact the lattice Conversely, taxol’s affinity at site *i* decreases if there is an excess of tau relative to taxol at site i or at the two neighboring sites *i* − 1 and *i* + 1 leading to the desorption of taxol from that site. Therefore, the floor-field effectively models the competition between tau and taxol on microtubules.

### 2.3 A static floor-field to model irregularities at random lattice sites

As observed in the experimental kymograph, tau envelopes can stop growing before reaching the ends of the microtubule. We hypothesize that this occurs due to irregularities in the lattice which could be due to a local chemical change or missing tubulin units. Indeed, lattice defects have been observed in several studies [25–28] and also modeled [29]. To account for these irregularities, we introduce a static floor-field in addition to the dynamic one. This static floor-field corresponds to changes in certain site properties that inhibit further envelope growth.

In our model, the coupling factor *h* is the key parameter that enables the envelopes to spread. Therefore, we naturally selected it as the variable whose alteration could prevent envelope growth at certain sites. When the coupling factor, *h*, is set to zero, there is no envelope formation as the adsorption of tau on a site is completely independent of the adjacent sites. To halt envelope growth, we reduce *h* to a very low value *h*^*t*^ at selected lattice sites. This reduction in coupling factor does not entirely prevent the envelope formation, which is consistent with the experimental results (Fig. 5). The sites with irregularities are randomly selected from a uniform distribution, with their number adjusted to produce envelope widths comparable to the experimental observations.

### 2.4 Hysteresis allows reduction in tau density in envelopes during envelope disassembly

In experiments where tau is kept in solution during and after envelope formation, the tau density in the envelopes stays constant. However, in experiments where tau is washed off from solution after envelope formation, the tau density within the envelopes decreases before the envelopes eventually disintegrate. This indicates that stable envelopes are allowed at lower densities in the envelope disintegration process than in the envelope formation process which can be explained by a hysteresis loop. Hysteresis is the property of a system’s evolution changing his path according to an external cause. In this case, the external factor is the absence or presence of tau in solution. The transition from a state with tau envelopes on a microtubule to a state without envelopes occurs at lower *δ*_*i*_ compared to the transition from no envelopes to the envelope formation on a microtubule. This shift allows for lower tau densities during envelope disintegration than during envelope formation. The transition between these two states is determined by the floor-field *H*_*i*_ and the hysteresis loop is constructed with two *H*_*i*_ shifted on the *δ*_*i*_ axis (Fig. 2). The amplitude of this shift, denoted as ∆*c*, determines the strength of the hysteresis.

## 3 Results

The goal of this study is to determine whether the tau-taxol competition can explain the envelope formation and disassembly observed in the *in vitro* experiments reported in Siahaan et al. [12]. Experimental data showed that when fluorescently labeled tau was added to taxol-stabilized microtubules, tau initially bound to the microtubule at a low density. This was followed by the nucleation of high-density tau patches, termed ‘tau envelopes’. These tau envelopes originated from diffraction-limited spots and grew at varying rates in both directions along the microtubule. In regions where tau does not form an envelope, referred to as the ‘surroundings,’ a lower density of diffusive tau still binds to the microtubule lattice.

To address our question, we use Monte Carlo simulations to replicate the experimental results *in silico*. Our analysis focuses on (i) the tau density within the envelopes and their surroundings, (ii) the fraction of the microtubule covered by envelopes, and (iii) the temporal evolution of the envelope boundary positions as recorded in the kymographs. In addition to the envelope formation, we also examine the envelope disassembly. The methodology employed to calibrate the model parameters is delineated in the Methods section. The optimal parameters, which successfully reproduce the experimental kymographs, tau density and coverage, are reported in Table 1. All rates correspond to the number of tau molecules per second on a given lattice site. For instance, a tau adsorption rate of 0.094 *s*^−1^ indicates that, on average, 0.094 tau molecules adsorb on each site per second. Our model does not account for concentration dependency, which is intrinsic to experimental measurements. Therefore, these quantities may not directly correspond to experimental units. The following sections compare the experimental measurements [12] with the simulation outcomes using optimized parameters.

**Table 1:** Table of optimized model parameters to match the experimental results. *h*^*t*^ is the value of the coupling factor on irregularities, *c*_*f*_ is the inflection point of *H*_*i*_ for the envelope formation and *c*_*d*_ is the inflection point of *H*_*i*_ for the envelope disassembly.

| Kinetic rates | Floor-field parameters |
| --- | --- |
| $\omega_a^\tau = 0.094 \text{ s}^{-1}$ (tau adsorption rate) | $\mu = 3$ (affinity modification factor) |
| $\omega_d^\tau = 0.43 \text{ s}^{-1}$ (tau desorption rate) | $h = 2$ (coupling factor) |
| $\omega_a^T = 0.2 \text{ s}^{-1}$ (taxol adsorption rate) | $h' = 0.2$ |
| $\omega_d^T = 0.3 \text{ s}^{-1}$ (taxol desorption rate) | $c_f = 8.14$ |
| $s = 50 \text{ s}^{-1}$ (tau diffusion rate) | $c_d = 3.5$ |
|  | 7% irregularities |

### 3.1 Tau-taxol competition is sufficient for tau envelope formation

#### Tau and taxol densities within envelopes and surroundings

We compare the simulated tau densities within envelopes and surroundings to the experimentally found values during envelope formation [12]. Fig. 3 shows the simulated tau density, with the parameter values listed in the Table 1 and the experimental results for 20 nM tau in solution [12]. From these findings we can make multiple observations. Firstly, we observe that the competition between tau and taxol is effectively represented by their respective densities: within regions of high tau density, taxol density approaches zero, whereas in surrounding areas, taxol density exceeds that of tau. This result is particularly consistent with experimental findings, which demonstrate a reduction in taxol density within the tau envelopes (see Fig. 1i,j in [13]). Secondly, we verify that we retrieve the Langmuir density given in Eq. (1) with the *intrinsic* rates for tau and taxol in surroundings, where the floor-field is zero. We retrieve the tau density of 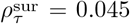 molecules per tubulin dimer as found in the experiments. Additionally, the tau density within the envelopes reaches the maximum density allowed by our coarse grain model 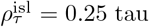 tau molecules per tubulin dimer, which aligns with the median value observed in the experiments. Finally, we note a discrepancy between the simulation and experimental results regarding the time when the first envelopes start to nucleate. In the experiment, envelope nucleation begins around *t* ≈ 100 seconds after tau is added to the solution, whereas in our simulations, envelopes appear within the first few seconds. However, it is important to clarify that the experimental densities are based on five microtubules only, while the theoretical results are averaged over a large number of simulated microtubules to provide a general result. Thus, the experimental result may not capture the full range of nucleation times, whereas even relatively rare nucleation times must be captured by the simulation result.

**Figure 3:**
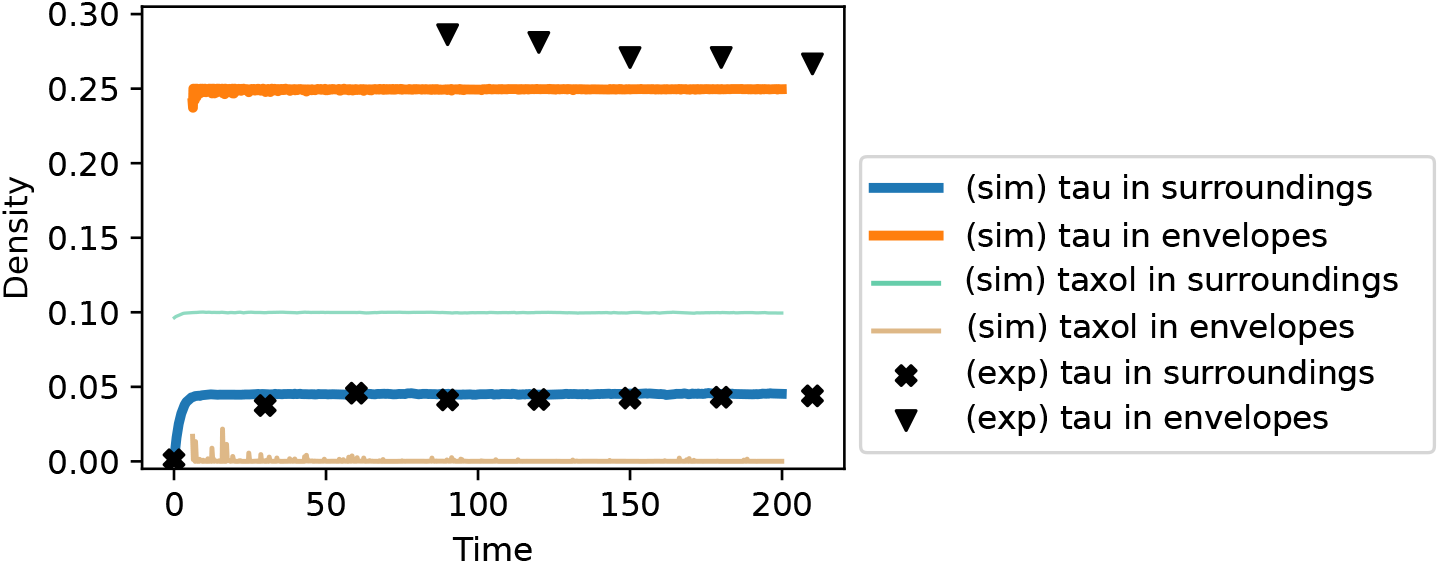
Densities obtained with the parameter values of Tab. 1 (*c* = *c*_*f*_) during envelope formation. The initial configuration is 20 taxols and 0 tau on each site. The results shown are the mean over 199 simulated microtubules with 100 sites. The black symbols present experimental time traces of the tau densities on taxol-stabilized microtubule following the addition of tau in solution (median over 5 microtubules), published in [12].

#### Fraction of microtubule covered by tau envelopes

The simulation results in Fig. 4 show a gradual increase in tau envelope coverage reaching the value of 10% at *t* = 200 *s* as observed in the experiment. However, in simulations the growth follows a linear curve, in contrast to the experimental observation. In Fig. 1e of [12] (replotted in Fig. 4), the coverage increases rapidly immediately after tau is introduced into the solution and then slows down progressively over time. We propose that this non-linear behavior arises from fluctuations in tau concentration in solution: upon tau injection, there is an initial burst of envelope nucleation, followed by a decline in nucleation as tau concentration diminishes due to imperfect conditions in the experimental setup. This behavior is not observed in simulations, as tau concentration is not incorporated into the model. Importantly, this increase in tau envelope coverage indicates that multiple envelopes nucleate within the first few seconds of the experiment, confirming that the late nucleation observed in the experimental density profile (Fig. 3) is not a universal occurrence.

**Figure 4:**
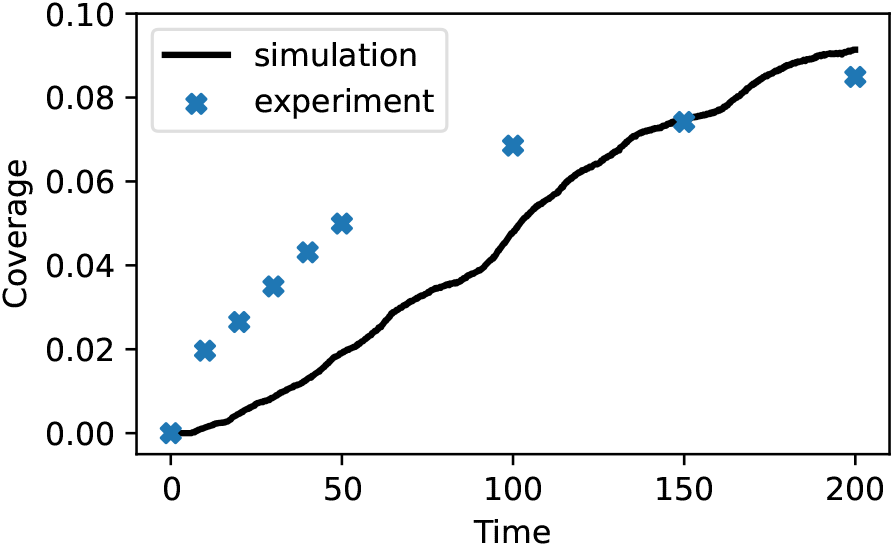
Fraction of microtubule length covered by tau envelopes obtained with the parameter values of Tab. 1 (*c* = *c*_*f*_) during envelope formation. The initial configuration is 20 taxols and 0 tau on each site. The results shown are the mean over 199 simulated microtubules of 100 sites. The experimental coverage of tau envelopes on taxol-stabilized microtubules following the addition of 20 nM tau in solution from [12], is also included, indicated by blue symbols (median over 131 microtubules).

#### Tau envelope formation kymograph

Fig. 5 compares a simulated kymograph with an experimental kymograph. The model successfully generates a large, stable envelope with a uniform tau density both within the envelopes and in the surroundings, as observed in the experimental data. Furthermore, the model allows for the merging of adjacent envelopes. However, a difference appears in the boundary produced by the envelope growing: in the experimental kymograph, this boundary is jagged, whereas it appears very regular in the simulation. This discrepancy could potentially be addressed by modifying the floor-field to varying its strength at the boundaries, a refinement that should be explored in future work.

**Figure 5:**
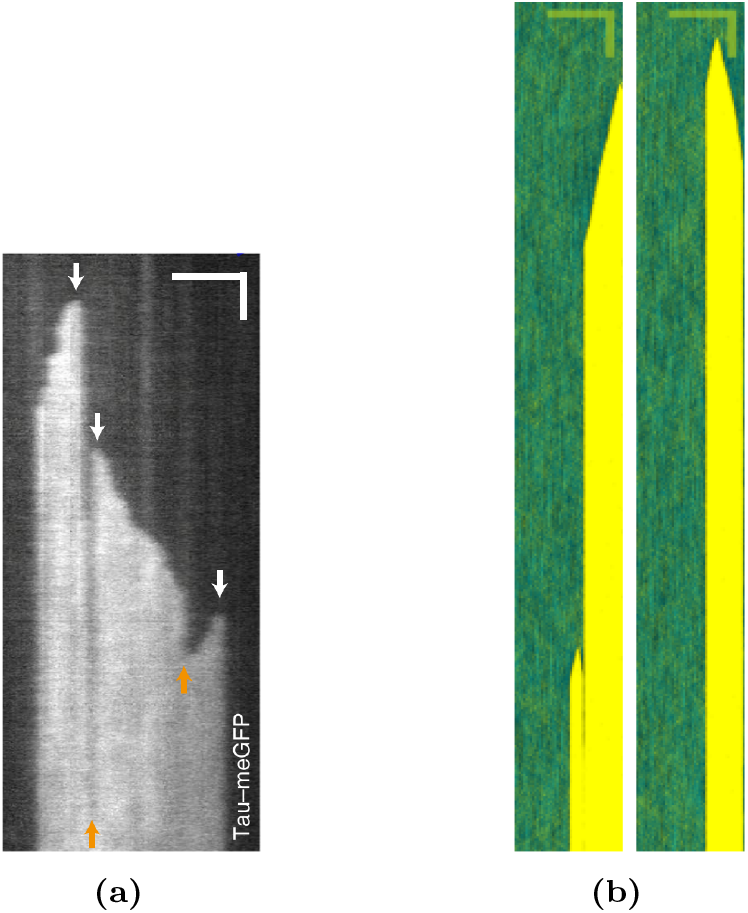
(a) Experimental kymograph showing the fluorescence signal of tau tagged with a fluorophore on a microtubule following the addition 20 nM tau. White arrows mark nucleation sites, while orange arrows indicate the merging of neighboring envelopes growing towards each other. Reproduced from [12]. (b) Two simulated kymographs of a microtubule 100 sites during envelope formation starting from the initial configuration of 20 taxols and 0 tau on each site. Tau molecules are represented in yellow whereas taxol molecules are represented in blue. Transparency allows seeing the superposition of the particles (tau on top). Scale bars: (horizontal) 2 *μm*, (vertical) 50 *s*.

### 3.2 Hysteresis allows for gradual envelope disassembly by a decrease in tau density

In the article [12] the kinetics of tau envelope disassembly, upon removal of tau from solution, was also investigated. In these experiments, microtubules were fully covered by tau envelopes by incubating the microtubules with a high concentration of tau molecules. After removing tau from solution, a gradual reduction in envelope coverage was observed.

To simulate the experiment, we take the initial condition of 13 tau and 0 taxol molecules on each site. To model the absence of tau in solution, we set the adsorption rate of tau to zero. While this is a simplification — since in the actual experiment, desorbed tau could potentially re-adsorb — it is a reasonable approximation, to not include a finite reservoir, for this first approach of modeling tau-taxol competition using a floor-field approach. As detailed in our modeling description, we introduced a hysteresis loop to capture the density decrease within envelopes. The simulation results aligned with experimental observations when the hysteresis parameter *c*_*d*_ was set to 3.5.

#### Tau density during envelope disassembly

We compare the simulated tau densities within envelopes and surroundings to the experimentally found values during envelope disassembly [13]. It should first be noted that our model assumes a tau molecule always covers four tubulin dimers, whereas in reality, a tau molecule can bind to 1, 2, 3, or 4 tubulin dimers. Consequently, a density greater than 0.25 tau molecules per tubulin dimer is not possible in our model. The primary objective here is to reproduce the overall behavior of the densities rather than the exact values.

As shown in Fig. 6, the simulation successfully captures the general behavior of the tau densities: the tau density within the envelopes decreases until *t* ≈ 200 *s*, followed by a slight increase until *t* ≈ 300 *s* and then decreases again. We will see by observing the kymograph that the non-monotonous density profile must be due to a combination of a boundary effect in which the tau molecules have the tendency to migrate near the boundary and the envelope fissuring. The main discrepancy is that the simulated envelopes do not persist beyond 500 seconds, unlike the experimental envelopes which last longer. To explain this discrepancy, we propose that the rebinding of tau after desorption slows down the overall disassembly of the envelope in the experiment, whereas tau cannot rebind in the simulation since the adsorption rate is set to zero. Moreover, this discrepancy is not universal, as some simulated envelopes disassemble much faster (within 100 seconds). Therefore, we can conclude that the simulation effectively captures the average behavior of envelope disassembly.

**Figure 6:**
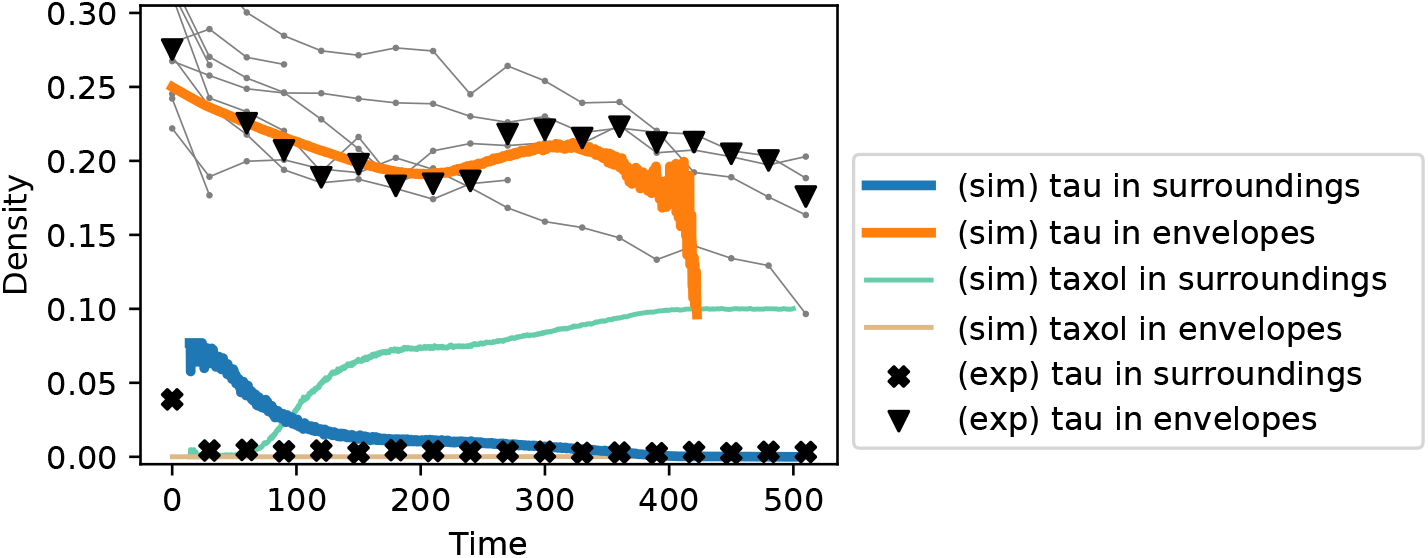
Densities obtained with the parameter values provided in Table 1 (*c* = *c*_*d*_) during envelope disassembly. The initial configuration is 13 tau molecules and no taxol at each site. Average over 99 simulated microtubule of 100 sites. The black symbols represent experimental time traces of the tau densities on taxol-stabilized microtubule following the removal of tau from solution (median over 9 microtubules), published in [12]. We also show tau densities in envelopes for individual microtubules (gray lines) to demonstrate that some envelopes disassemble in less than 500 seconds after removal of tau from solution.

Regarding the tau density in the surrounding areas, the simulation shows a rapid decrease, albeit slightly slower than observed in the experiments. This discrepancy arises from the simplistic method used to differentiate between envelope and surrounding areas in the simulation. In the model, a site is considered part of an envelope if it has at least four tau molecules adsorbed on it and its neighboring site; all other sites are categorized as surrounding. Initially, the lattice is fully covered by an envelope, so the density in the surrounding areas may be overestimated as these regions are only close to the envelope boundaries. However, the time required for the density to drop from the initial 0.045 tau molecules per tubulin dimer to 0.001 tau molecules per tubulin dimer with 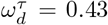 is given by Eq. (9) and is only 9 seconds, which is consistent with experimental measurements showing that tau density reaches zero within 30 seconds in the regions surrounding envelopes.

Finally, we can reiterate that the tau-taxol competition is effectively represented by the densities: as the tau density in surroundings decreases to zero, the taxol density increases substantially.

#### Tau envelope disassembly Kymograph

Fig. 7 compares a simulated kymograph with an experimental kymograph of tau desorption from a microtubule upon removal of tau from solution. We can see that the simulation closely matches the experimental observation of envelope disassembly over time. However, it is noteworthy that the simulated kymograph shows a lower tau density near the center of the envelope and throughout the bulk of the two legs compared to the envelope boundaries, which denotes the presence of a boundary effect. This boundary effect is highlighted by the tau density profile in the Supplementary Fig. 1. Additional simulations that did not exhibit this boundary effect also lacked the local density maximum after the envelope fissure (Supplementary Fig. 2). This finding indicates that the boundary effect likely contributes to the observed local maximum in density following envelope fissuring. As the envelope splits, the boundary length increases, leading to a higher proportion of high-density regions, which, in turn, causes an overall increase in mean density.

**Figure 7:**
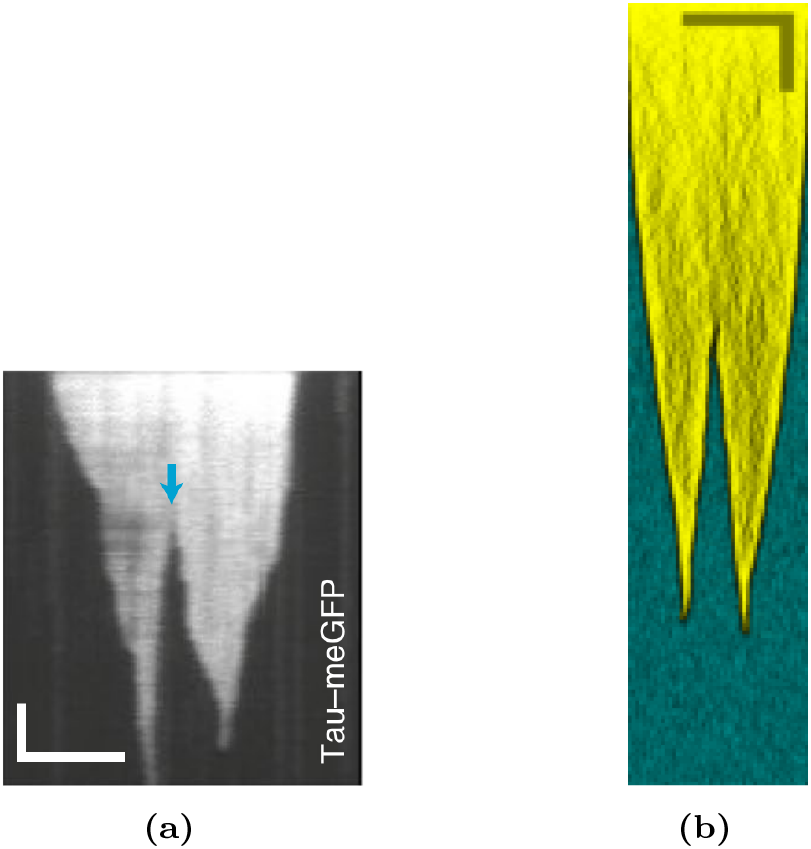
(a) Experimental kymograph showing the fluorescence signal of tau tagged with a fluorophore on a microtubule following the removal of tau from solution. The blue arrow indicates a fissuring event. Reproduced from [12]. (b) Simulated kymograph of a microtubule 100 sites starting from the initial configuration of 0 taxol and 13 taus on each site. Tau molecules are represented in yellow whereas taxol molecules are represented in blue. Transparency allows seeing the superposition of the particles (tau on top). Scale bars: (horizontal) 2 *μm*, (vertical) 50 *s*.

## 4 Conclusion & discussion

In summary, we have demonstrated that a simple floor-field model of tau-taxol competition on a lattice with irregularities and a hysteresis loop can effectively reproduce the envelope formation and disassembly, along with their distinct features, as observed in experimental studies [11–13]. Taxol is the molecule used to stabilize microtubules in experiments. The tau-taxol competition is modeled by considering the lattice’s varying attractiveness: as tau compacts the lattice, it becomes more attractive to tau but less so to taxol; conversely, as taxol expands the lattice, it becomes more attractive to taxol but less so to tau. Specifically, we hypothesize that the transversal compaction increases the affinity for new taus on the neighboring sites of the adjacent protofilaments whereas the longitudinal compaction increases the affinity for tau on the neighboring sites on the same protofilament of the site bound to taus.

It has been suggested that tau-tau interaction play a key role in tau envelope formation [12]. However, previous work attempted to explain this phenomenon through tau-tau interaction but failed to validate the hypothesis [14]. The main difference of the floor-field model with modeling a biochemical tau-tau interaction is that the adsorption rate of tau is increased by the tau already adsorbed on the lattice, but taus adsorbed on the lattice do not attract each other. This allows to obtain a non-negligible density of tau in the surroundings (outside the envelope) whereas a tau-tau interaction would produce envelopes absorbing all the tau in the surroundings.

Our model successfully replicates the formation of stable envelopes with sharp boundaries, which halt growth at certain ‘irregularities’ of the lattice. The consideration of irregularities on the microtubule lattice is consistent with the observation of missing tubulins within a protofilament [29], and it is reasonable to assume that this could be the cause of the reduction in the propagation effect of lattice compaction. Additionally, envelopes that are close to each other can merge if their growth was previously stopped by an irregularity, mirroring experimental observations. We also capture the experimentally observed characteristics of envelope disassembly, which has been described as showing cohesiveness [12]. In our simulations, when tau can no longer adsorb onto the lattice, the envelope shrinks and eventually splits into a limited number of parts, while tau in the surroundings desorbs more rapidly. This demonstrates a lower propensity for tau desorption within the envelope compared to the surrounding areas. However, our model indicates that tau-tau interactions are not essential for this cohesiveness. Instead, the stronger affinity of tau for a compacted lattice, compared to an expanded one, is sufficient to explain the apparent cohesiveness of the tau envelope. Therefore, we conclude that the tau-taxol competition, driven by the lattice’s expansion and compaction, is enough to account for the tau envelopes observed on taxol-stabilized microtubule.

Our model introduces new elements that have yet to be experimentally observed. Future work should aim to provide experimental evidence for these elements. Key questions include: What are these ‘irregularities’ along the microtubule lattice and how do they halt envelope growth? What underlies the hysteresis loop? How does tau compact the microtubule lattice? Regarding the last question, it is noteworthy that Tan et al. [11] demonstrated that the projection domains flanking tau’s four microtubule-binding domains are necessary for the formation of the tau envelope. As we have shown that the lattice compaction by tau leads to the tau envelopes formation, we hypothesize that the projection domains are required for lattice compaction. Additionally, there are opportunities to further refine the model. Incorporating tau-tau interactions could provide insights into how these interactions influence tau-taxol competition and determine the threshold at which the model remains valid. We could also enhance the model by including minor inhomogeneities in the floor-field, which may yield more variable envelope growth and account for the different growth velocities observed in experimental kymographs. Moreover, exploring the effect of tau phosphorylation on envelope stability, as reported in recent studies [30], would be a valuable extension.

The model developed for tau envelope formation and disassembly also holds significance in the field of theoretical physics. It can be described as a Symmetric Simple Exclusion Process (SSEP) with Langmuir kinetics and a floor-field. The SSEP is a well-studied model in the context of one-dimensional lattice gases [31–35]. Thus, the analytical study of this model will not only deepen our understanding of the physical processes underlying tau envelope behavior, but also contribute to the broader field of lattice gas models.

## 5 Methods

### 5.1 Extracting quantities from experimental data

In the envelope surroundings, the floor-field has no strength since *H*_*i*_ = 0. Consequently, the tau and taxol densities are given by the Langmuir density (Eq. (1)). Although the taxol density in surroundings is not known, the measured tau density from the Fig. 1d of [12] is approximately 0.045 tau per tubulin dimer in surroundings, which provides the constraint:

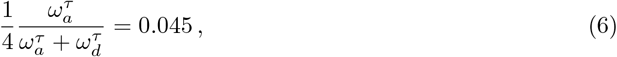

on the adsorption and desorption rates of tau. Here the factor of four accounts for the coarse graining of the microtubule lattice, where one site corresponds to 4 tubulin dimers.

From the measure of the tau envelope disassembly density we can infer the desorption rate of tau. In the disassembly experiments, tau molecules are removed from the solution, which we simulate by setting 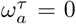. However, the tau density in the surroundings cannot directly provide 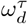 since the density is already almost zero at the first measurement point after the initial time. This means we cannot determine if the density had already approached zero before the first measurement, potentially underestimating the slope. There are not enough data points to accurately measure the slope to zero. We can only conclude that 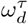 must be equal to or greater than the value suggested by the slope, which corresponds to 0.08 tau molecules per tubulin dimer. Therefore, we focus on the tau density within the envelope, where the floor-field modifies the rates (*H*_*i*_ *>* 0). To estimate 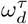 with the density decrease inside the envelope, we consider the effective desorption rate

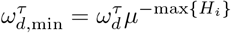

where max*{H*_*i*_*}* = 1 + 2*h* is the maximum value of the floor-field for a given coupling constant *h*. The equation of the tau density, in its site-independent form and when 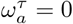 is

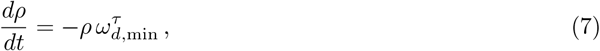

which describes the evolution of the tau density over time. Solving this differential equation, we obtain the tau density as a function of time:

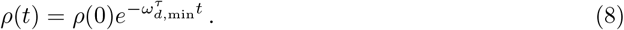

We can then estimate 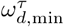 from the density measurement of Fig. 1h of [12] with the formula

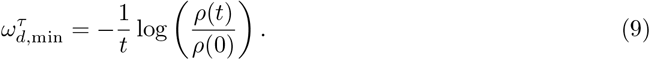

The simulated densities during envelope disassembly, as reported in the Fig. 6, show that the non-monotonic variation in density within the envelope should not be attributed to mere statistical fluctuations. To determine 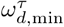, we measured the slope between the initial time point and the local minimum before the density increases again. Using the points *ρ*(*t* = 0) = 0.276 and *ρ*(*t* = 200) = 0.19, we estimate 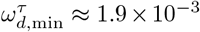. The effective desorption rate 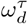 was then be adjusted based on the selected coupling constant *h* and the affinity modification factor *μ*.

### 5.2 Simulation specifications

Simulations are performed using the Gillespie algorithm for continuous time Monte Carlo [36, 37].

#### Envelope and surrounding differentiation

During envelope formation, a site is considered part of an envelope if it has at least eleven tau proteins adsorbed on it and its neighboring sites. During envelope disassembly, the threshold is lowered to four taus since the density inside the envelope decreases significantly, especially far from the envelope boundaries.

### 5.3 Adjusting trials

To accurately replicate the observed features in experiments, multiple simulation trials were necessary, as the model parameters influenced various aspects of the results. The effects of each floor-field parameter are summarized in Table 2.

**Table 2:** Main influences of the floor-field parameters on the tau envelope features. The positive (an increase) or negative (a decrease) influences are represented with arrows.

| Floor-field parameters | Features of the tau envelopes |
| --- | --- |
| $\mu$ | coverage (↗), tau density in envelopes (↗) |
| $h$ | spreading speed (↗), disassembly speed (↘) |
| $h'$ | envelope merging (↗) |
| $c_f$ | coverage (↘), tau density in envelopes during assembly (↗) |
| $c_d$ | tau density in envelopes during disassembly (↗) |
| Nbr. irregularities | envelope length (↘), coverage (↘) |

#### Taxol density

Currently, there are no direct measurements of taxol density or kinetic rates available. However, it is crucial that the taxol density is kept low enough to prevent the formation of taxol envelopes. Based on this, we selected a taxol density of 0.4 taxol molecules per tubulin dimer.

#### Irregularities and envelope length

In our simulations, the only parameter influencing the length of the tau envelope is the proportion of lattice irregularities. To match the envelope length observed in experiments during envelope formation, we then adjusted the proportion of irregularities. Simulating a lattice with 100 sites, corresponding to a small microtubule of 3.2 *μm*, we found that a 5% irregularity ratio yielded an envelope length similar to the experimental kymograph shown in Fig. 7a. Accounting for the microtubule boundaries, the effective irregularity ratio was determined to be 7%. This implies that similar results must be obtained in simulations for longer microtubules using a 7% irregularity ratio.

#### Affinity modification factor *μ*

The affinity modification factor *μ* controls the strength of the floor-field, thereby affecting the nucleation rate of tau envelopes. As *μ* increases, the effective rates change rapidly, facilitating the transition from the surrounding phase to the envelope phase. However, *μ* must not be set too high in order to retrieve the low proportion of microtubule covered by envelopes measured in the experiments. On the other hand, the tau density within the envelopes increases with *μ* and we aimed to match the experimental tau density of approximately 0.25 tau molecules per tubulin dimer, which is the maximum density allowed by our coarse-graining model (where a tau molecule covers four tubulin dimers). Thus, *μ* must be carefully balanced—sufficiently high to achieve the correct envelope density, but not so high as to overestimate microtubule coverage.

#### Coupling factors (*h, h*^′^)

The coupling factor *h* also influences the floor-field strength by increasing *H*_*i*_, but its primary effect is on the speed of envelope growth. We adjusted *h* to match the slope of the envelope growth as observed in the experimental kymographs. On sites with irregularities, the coupling factor is adjusted to *h*^′^, which must be low enough to halt envelope growth but high enough to allow adjacent envelopes to merge even with an irregularity in the middle.

#### Floor-field function centers (*c*_*f*_ and *c*_*d*_)

The center of the floor-field function *H*_*i*_ for the transition from surrounding to envelope, called *c*_*f*_, determines for which difference between the tau proteins and the taxol molecules *δ*_*i*_, the transition from the surrounding state to the envelope state begins to occur. Therefore, it tunes the nucleation rate, and then the coverage, by being more or less close to the stationary tau-taxol difference 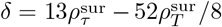 in surroundings. The *δ*_*i*_ at the right of the transition (higher *δ*_*i*_, see Fig. 2) corresponds the tau-taxol difference in the envelopes and then to the tau and taxol densities allowed within the envelopes. Consequently, *c*_*f*_ must be high enough to retrieve the low coverage and the high homogeneous density in the envelopes.

Inversely, the center of the floor-field function *H*_*i*_ for the transition from envelope to surrounding, called *c*_*d*_ determines for which difference between the tau proteins and the taxol molecules *δ*_*i*_, the transition from the envelope state to the surrounding state occurs. By setting *c*_*d*_ *< c*_*f*_, we allow the tau density decreasing in envelopes as it is observed in experiments. The value of *c*_*d*_ is tuned in order to retrieve the minimal density of 0.19 tau per tubulin dimer at *t* = 200 seconds after removing tau molecules from the solution in the experiment.

#### Diffusion Rate

The diffusion rate of tau molecules must be relatively high to account for the homogeneous tau densities in the envelope surroundings observed in the experimental kymographs. This high diffusion rate ensures that tau molecules spread evenly across the microtubule lattice.

## Supporting Information

Fig. S1. Boundary effect. Fig. S2. No boundary effect.

## Acknowledgments

The authors wish to thank Zdenek Lansky for helpful discussions, Reza Shaebani and Adam Wysocki for their support.

## Author Contributions

Conceptualization: Ludger Santen, Carole Chevalier

Investigation: Ludger Santen, Carole Chevalier Supervision: Ludger Santen

Visualization: Ludger Santen, Carole Chevalier

Writing – Original Draft Preparation: Ludger Santen, Valerie Siahaan, Rahul Grover, Stefan Diez, Carole Chevalier

