## Supplementary material for "A floor-field model of tau-taxol competition explains tau envelope dynamics on taxol-stabilized microtubules": Fig. S1.

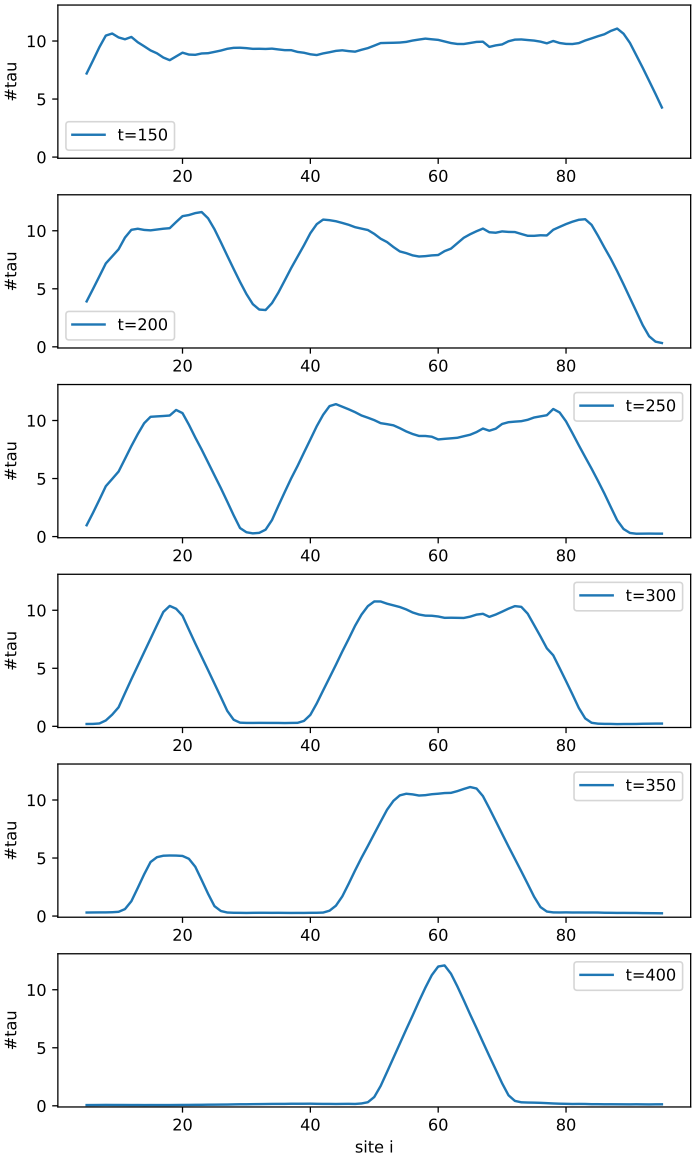


Rolling window. $\delta t = 12\,s,\ \delta l = 10$ sites.

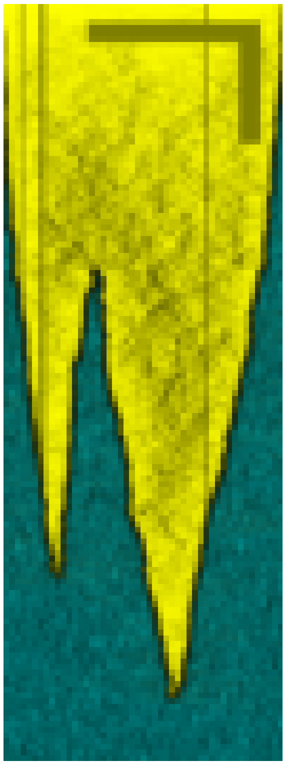

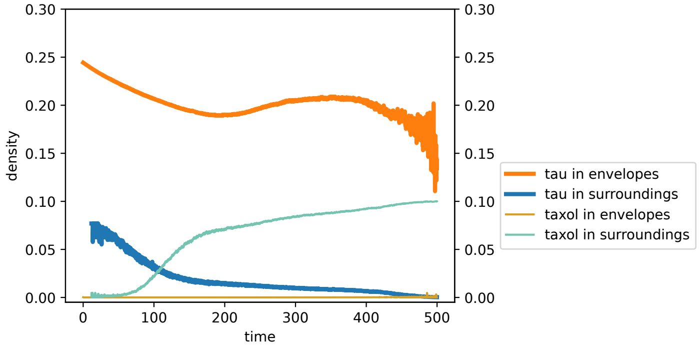


Fig. S1: **Boundary effect.** (left) Local number of tau molecules at different time of the simulation. (right top) Kymograph. Tau molecules are represented in yellow whereas taxol molecules are represented in blue. Scale bars: (vertical) time: 80 $s$, (horizontal) length: 2 $\mu m$. (right bottom) Tau and taxol densities. Average over 100 simulated microtubule. Parameters: $L = 100$ sites, $\omega_a^\tau = 0\ s^{-1}$, $\omega_d^\tau = 0.15\ s^{-1}$, $\omega_a^T = 0.02\ s^{-1}$, $\omega_d^T = 0.03\ s^{-1}$, $s = 75\ s^{-1}$, $\mu = 2.4$, $h = 2$, $c_d = 3$. Initial configuration: 0 taxol and 13 taus on each site. In the bigger leg, we see clearly that the density is bigger at the boundaries.
